# A feline model of human LDLR-related atherosclerosis

**DOI:** 10.1101/2024.12.04.626782

**Authors:** Marjo K Hytönen, Veera Karkamo, Sruthi Hundi, Niina Airas, Maria Kaukonen, Antti Sukura, Leslie A Lyons, Heidi Anderson, Ilona Kareinen, Hannes Lohi

## Abstract

**Background:** Atherosclerosis, a chronic inflammatory vascular disease driven by the accumulation of LDL-derived cholesterol on arterial walls, is the leading cause of mortality worldwide but is rare in animals. We recently identified spontaneous atherosclerosis in the Korat cat breed, characterized by severe hypercholesterolemia and clinical signs of congestive heart failure, ultimately leading to death. Histopathological examination revealed lesions similar to those observed in human atherosclerosis. Given the close genetic relationship among affected cats, we hypothesized a genetic basis for the condition.

**Methods:** We expanded our sample recruitment and employed whole genome sequencing to identify genetic variants associated with the condition.

**Results:** We identified a homozygous XM_003981898.6:c.2406G>A variant specific to the cases in the *LDLR* gene. This variant is predicted to result in a premature stop codon, XP_003981947.3:p.Trp758*, leading to a truncated LDLR protein that lacks the last 108 amino acids, including the transmembrane and intracellular C-terminal domains. Genotyping this *LDLR* variant in an additional cohort of 309 Korat cats confirmed its segregation and revealed new affected cats for clinical follow-up. In silico analyses demonstrated that the identified variant appears optimal for gene-editing-based therapeutics.

**Conclusions:** This is the first report of a spontaneous atherosclerosis animal model with an *LDLR* variant, the most common gene associated with familial hypercholesterolemia in humans. Given that *PCSK9*, another known hypercholesterolemia gene, has been lost in many mammalian genomes, including cats, our study provides an exciting double knockout model for human atherosclerosis. The affected Korats may also serve as a valuable model for DNA base editing therapeutics.

## Introduction

Atherosclerosis is a chronic inflammatory vascular disease initiated by the accumulation of LDL-derived cholesterol on the arterial inner surface. Human familial hypercholesterolemia is the most potent risk factor for atherosclerosis and the three most frequently associated genes are *low density lipoprotein receptor* (*LDLR), apolipoprotein B* (*APOB)*, and *proprotein convertase subtilisin/kexin type 9* (*PCSK9)*, where *LDLR* being the most common with over 2000 pathogenic variants [1–4]. However, natural animal models with *LDLR* or other gene defects have not been reported. Interestingly, some mammalian species, including cats, have lost *PCSK9* during evolution.

A spontaneous atherosclerosis with severe hypercholesterolemia, clinical signs of lethal congestive heart failure, and histopathology resembling human disease, was recently described in cats [5]. We describe here the genetic cause of feline atherosclerosis: a loss-of-function variant in *LDLR*.

## Material and Methods

The study cohort consisted of 309 blood and tissue samples from European and US Korat breed cats, including two published atherosclerosis cases [5].

We conducted whole-genome sequencing (WGS) on a trio of the Korat cats (one affected and two unaffected parents) (Fig. 1A) at Novogene (HK) Company Ltd using the Illumina HiSeqX platform with ∼30x coverage. We used the DRAGEN pipeline from the Illumina DRAGEN Complete Suite v3.9.5 to map and align the raw data to the Felis_catus_9.0 genome assembly [6] and for joint genotyping of this data to create a multi-VCF. The genomic and nucleotide positions correspond to the latest feline genome assembly F.catus_Fca126_mat1.0 (GCF_018350175.1) and *Felis catus* Annotation Release 105. We validated the candidate variant identified from the WGS analysis by PCR and Sanger sequencing using the following primers: 5’-TTCTTTCAGAGCCCGAACCT and 3’-GAAAGTGCAGGGGTGAAAGG. To confirm the clinical phenotype, we invited one cat for a clinical follow-up study to measure cholesterol levels and performed a post-mortem examination for one cat that died during the study. Patients or the public were not involved in the design, or conduct, or reporting, or dissemination plans of our research.

**Figure 1.**
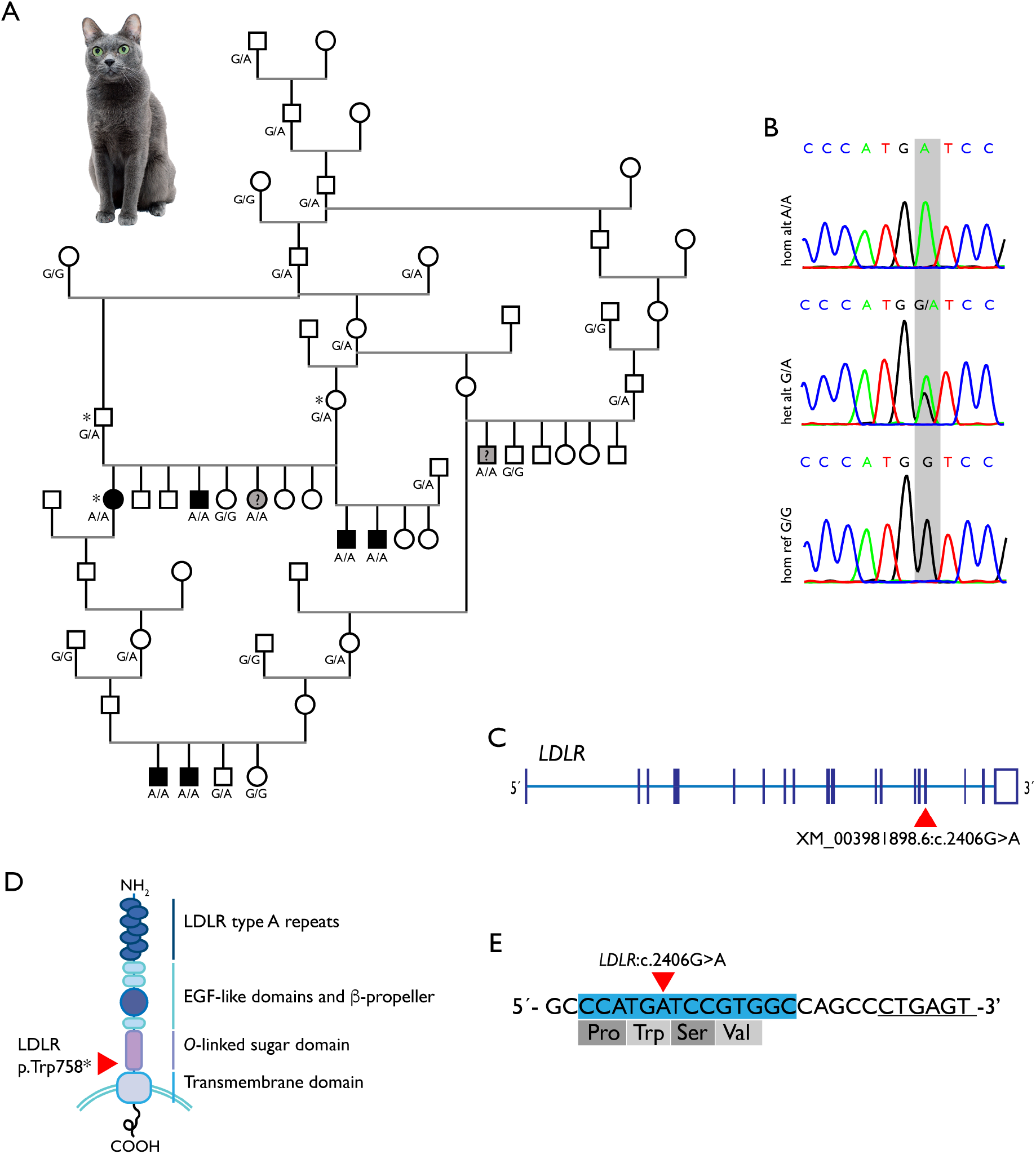
A. An extended pedigree for Korat cats with LDLR-related hypercholesterolemia. The affected and *LDLR* variant homozygous cats marked as black and *LDLR* variant homozygous without confirmed clinical phenotype marked as grey and question mark (?). The *LDLR* variant genotypes of the cats available for the study are indicated below the symbols. The whole genome sequenced cats indicated by asterisk (*). Korat cat photo by Marko Lumikangas. B. Chromatograms indicating the genotypes of an affected cat homozygous for the alternative allele (hom alt, A/A), heterozygous (het alt, G/A) and wild-type (hom ref, G/G) cats. C. A schematic exon-intron structure of the feline *LDLR* with the variant site (red arrow head) indicated. D. A schematic figure of the LDLR protein domains based on human protein structure. The LDLR c.2,2406G>A variant is predicted to result in a truncated protein (p.Trp758*, red arrow head), if translated, lacking the C-terminal transmembrane and intracellular domains. E. A highly potential DNA base editing site indicated around the mutation site (red arrow head) in *LDLR*. Base editing window shown in blue, the PAM site underscored.

## Results

We identified variants in the candidate genes *LDLR* and *APOB* through WGS. *PCKS9* has been lost in many mammalian genomes, including cats [7], and could not be tested. Recessive filtering revealed a homozygous loss-of-function variant specific to the case in *LDLR* on a genomic position A2:8410638G>A (F.catus_Fca126_mat1.0) (Fig. 1B-D). The XM_003981898.6:c.2406G>A variant is predicted to result in a premature stop codon, XP_003981947.3:p.Trp758*, and a truncated LDLR protein, removing the last 108 amino acids including the transmembrane and intracellular C-terminal domains (Fig. 1C, D). Notably, this variant was absent in 340 feline genomes of different breeds in the 99 Lives Cat Genome Consortium dataset [6].

The genotyping of 309 Korat cats for the stop-gain variant identified 239 wildtype, 62 heterozygous, and 8 homozygous individuals. Of the homozygous cats, three were diagnosed with atherosclerosis at necropsy, two with hypercholesterolemia, one succumbed at 7 years of age to a hind limb thrombus. Two of the cats remained without confirmed clinical phenotype as one had died due to kidney failure at 15 years of age, and one at 5 months of age for an unknown reason. Total cholesterol levels, available for three of the homozygous cats, were markedly elevated (12.38–25 mmol/L). Owners did not report any signs of cardiovascular disease in the heterozygous cats; however, a comprehensive lipid profiling is warranted and ongoing to better understand genotype-phenotype correlations.

Our *in silico* analysis indicates that the feline *LDLR* variant is an optimal candidate for CRISPR-Cas9-mediated DNA base editing [8]. As a G-to-A variant, it can be corrected back to the wild-type sequence using an adenine base editor (ABE). Additionally, a suitable *S. aureus* 5’-NNGRRT-3’ PAM site is located downstream of the variant, positioning the mutation within the editing window of the construct. The only predicted bystander edit would result in a silent p.Pro757Pro alteration (Fig. 1E).

## Discussion

In this study, we introduce a valuable naturally occurring animal model for familial hypercholesterolemia (FH) in humans caused by a defective LDLR. Homozygous cats with elevated cholesterol levels developed severe atherosclerotic lesions and exhibited signs of cardiovascular disease, yet notably survived into middle and advanced age.

In the affected cats, the atherosclerotic lesions advance to complex fibroatheromas with massive cholesterol accumulation, disruptive lipid cores and calcium deposits. Unlike other animal models, we provided evidence that endothelial disruption may predispose these cats to thrombosis [5]. Similar to humans, cats with likely dysfunctional LDLR experience a silent progression of atherosclerosis over several years, with clinical signs appearing only at advanced stages. In this feline model, the most advanced lesions occur in large elastic arteries, but coronary artery lesions, which can lead to ischemia and myocardial damage, are also a notable feature [5].

Although formal confirmation of the absence of LDLR protein in Korats is still pending, our feline model likely represents an intriguing double knockout of the two common FH genes, *LDLR* and *PCSK9*. PCSK9 typically regulates LDLR by promoting its degradation in the liver, thus reducing the liver’s ability to clear LDL cholesterol from the bloodstream. However, the *PCSK9* gene was lost early in Carnivora evolution, before the split between Caniformia and Feliformia [9]. The absence of PCSK9 in cats may promote greater recycling of LDL receptors, improving cholesterol clearance and contributing to their general resistance to atherosclerosis. Despite their high-protein, high-fat diet as obligate carnivores [10], cats appear to efficiently regulate cholesterol levels, with arterial disease rarely reported.

In the Korat cats with the homozygous stop-gain variant in *LDLR*, elevated cholesterol levels indicate impaired LDLR function. The lack of PCSK9 might help to maintain adequate cholesterol level in the heterozygous cats. This could explain why LDLR-related atherosclerosis appears to be a recessive condition in cats, unlike the predominantly dominant condition seen in humans. Ongoing lipid profiling of these heterozygous cats may provide further insight into this recessive pattern.

The feline *LDLR* variant appears highly accessible for base editing because of its compatibility with adenine base editors, its ideal placement within the editing window due to a downstream S. aureus PAM site, and the lack of predicted amino acid-altering bystander edits. Thus, these cats could provide a promising preclinical model for FH research and potential gene-editing therapeutics, and we have ongoing cell culture experiments to test the efficacy and safety of this approach.

